# An outlook of plants alliance identified on land and coastland, Southeastern Europe

**DOI:** 10.1101/2025.08.28.672895

**Authors:** Kuenda Laze

**Affiliations:** Polytechnic University of Tirana, Faculty of Civil Engineering

**Keywords:** Albania, Bosnia and Herzegovina, Croatia, Montenegro, Slovenia, land, plants

## Abstract

Land and coastland vegetation is diverse in Southeastern Europe. Plant species are identified and monitored. Yet, there are plant species not fully observed in Southeastern Europe. This study investigated verified and uncertain plant (herb, shrub, tree) alliances occurrences data extracted from Database of European Vegetation, Habitats and Flora until the year 2024, for Albania, Bosnia and Herzegovina, Croatia, Montenegro and Slovenia of Southeastern Europe using descriptive statistics. Verified herb, shrub and tree alliance occurrences were respectively 64.5 percent, 56.3 percent and 21.7 percent, and uncertain herb, shrub and tree alliance occurrences on land were, on average, respectively, 70.5 percent, 13.9 percent and 15.4 percent in the five Southeastern European countries. The uncertain herb alliance percentage tended to be higher compared to the uncertain shrub and tree alliance percentage in the five countries.

## Introduction

Vegetation contributes to carbon storage, and biodiversity ecosystem services. Monitoring of short-term changes and those long-term in vegetation was observed by collecting vegetation data using field sampling and remote sensing (Cupertino *et al*., 2024). Overall, field work, expert knowledge and remote sensing are tools used today for data gathering on vegetation.

The complete documentation of biodiversity at (vegetation) species and (vegetation) habitat level is of great importance (Chytrý *et al*., 2024) for biodiversity, nature conservation, habitat monitoring (and or habitat restoration) and research (Novák *et al*., 2023). Detailed information about the distribution of vegetation types and vegetation habitat types is important for basic ecological and biogeographical research (Preislerová *et al*., 2022). Preislerová *et al*., (2024) provided the first comprehensive compilation of standardized data on European vegetation types, based on expert knowledge and literature knowledge, providing standardized variables and attributes making it possible to query the database using a combination of various criteria to identify particular sets of alliances with similar structure, ecology and biogeography. This is important for regular repetitive vegetation habitat monitoring at various spatial scales (Lausch *et al*., 2016) e.g., at regional level, to better understand the changes in vegetation happening and the missing vegetation data at regional level, while considering ongoing climate change effects on vegetation and biodiversity decline in vegetation.

Here, plant alliances were used to understand the vegetation data collection at regional level, using descriptive statistics of vegetation alliances data retrieved from Database of European Vegetation, Habitats and Flora (Preislerová *et al*., (2024)) until the year 2024, for neighboring countries of Albania, Bosnia and Herzegovina, Croatia, Montenegro and Slovenia in Southeastern Europe. This study focused on identifying (i) verified and uncertain plant alliance (dominating life form of herb, shrub and tree) occurrences on land and coastland of five countries of Albania, Bosnia and Herzegovina, Croatia, Montenegro, Slovenia (ii) differences of verified and uncertain plant alliances between land and coastland and between countries of Albania, Bosnia and Herzegovina, Croatia, Montenegro, and Slovenia.

## Method

Plant alliances were extracted from Database of European Vegetation, Habitats and Flora (floraveg.eu) until the year 2024, being provided as an excel file version 2 (2024-06-12), matching the updated vegetation classification of EuroVegChecklist version 3 (2024-06-12) containing the highest vegetation cover in at least one of the four main vegetation layers (i.e., tree, shrub, herb or moss layer) as verified plant occurrence numbered “1” and uncertain plant occurrence texted “U” (Preislerová *et al*., 2024). Verified and uncertain plant occurrences were respectively extracted for plants on land and coastland for Albania, Bosnia and Herzegovina, Croatia, Montenegro and Slovenia resulting in ten columns. Verified or uncertain herb, shrub or tree alliance occurrences were separately calculated into 1-“trees and herbs” or “trees, shrubs and herbs” or “trees and shrubs”, or “trees” i.e., dominating life form of trees and shrubs and or herbs and of trees, 2 - “shrubs and herbs” or “shrubs” that is dominating life forms of “shrubs and herbs” or “shrubs”, and 3-“herbs” (dominating herbs and low vegetation as lichens, moss). Empty cells presenting absence of plant occurrences (Preislerová et al., 2024) were not considered in this data analysis. Information about plant alliance data collection and processing can be found at Preislerová *et al*., (2024).

The verified and uncertain plant occurrences were checked with 1- the total number of verified or uncertain plant alliance occurrences of the dataset, and with 2- the number of verified or uncertain tree, shrub and herb alliance occurrences on land and coastland of Albania, Bosnia and Herzegovina, Croatia, Montenegro, Slovenia (not calculated and provided in the dataset).

Descriptive statistics method was applied to plant alliance data analysis. The questions were as follows:1-What are the verified and uncertain plant alliance occurrences in Albania, Bosnia and Herzegovina, Croatia, Montenegro, Slovenia? 2-Are there differences in terms of verified and uncertain plant alliance occurrences between land and coastland amongst the five countries of the study region?

## Results

### Herbs, shrubs and trees

The number of verified plant alliance occurrences were higher on land than on coastland for the five countries. The lowest uncertain unverified plant alliance occurrences were in Slovenia (24) and Croatia (30) and the highest number of uncertain plant alliance occurrences was on land in Albania (77). The highest number of verified plant alliance vegetation occurrences were in Croatia (15), Albania (14) and Montenegro (13), on coastland (Table 1).

**Table 1.** The total number of verified and uncertain plant alliance occurrences on land and coastland in Albania, Bosnia and Herzegovina, Croatia, Montenegro and Slovenia (source: floraveg.eu).

| Land | Verified plant alliance occurrences | Uncertain plant alliance occurrences | Coastland | Verified plant alliance occurrences | Uncertain plant alliance occurrences |
| --- | --- | --- | --- | --- | --- |
| Albania | 133 | 77 | Albania | 14 | 4 |
| Bosnia and Herzegovina | 162 | 40 | Bosnia and Herzegovina | 4 | 2 |
| Croatia | 172 | 30 | Croatia | 15 | 3 |
| Montenegro | 146 | 59 | Montenegro | 13 | 4 |
| Slovenia | 165 | 24 | Slovenia | 7 | 2 |

Verified herb, shrub and tree alliance occurrences on land, on average, were respectively 64.5 percent, 56.3 percent and 21.7 percent, and uncertain herb, shrub and tree alliance occurrences were, respectively, 70.5 percent, 13.9 percent and 15.4 percent in the five Southeastern European countries. On coastland, verified herb, shrub and tree alliance occurrences were, respectively, on average, 82.0 percent, 10.1 percent and 7.87 percent, and uncertain herb, shrub and tree alliance occurrences were 46.7 percent, 33.3 percent and 20 percent.

The highest uncertain herb alliance percentage and uncertain tree alliance percentage on land was respectively in Montenegro (83 percent), Bosnia and Herzegovina (77 percent), Albania (74 percent) and in Croatia (23 percent), Figure 1. Overall, the highest percentage of verified and uncertain plant alliance occurrences were herbs in all countries. Yet, there were differences between verified and uncertain plant alliance occurrences on land and coastland and amongst the five study countries, e.g., the percentage of uncertain herb alliance occurrences were higher in Montenegro, Bosnia and Herzegovina and Albania than in Croatia and Slovenia (Figure 1).

**Figure 1.**
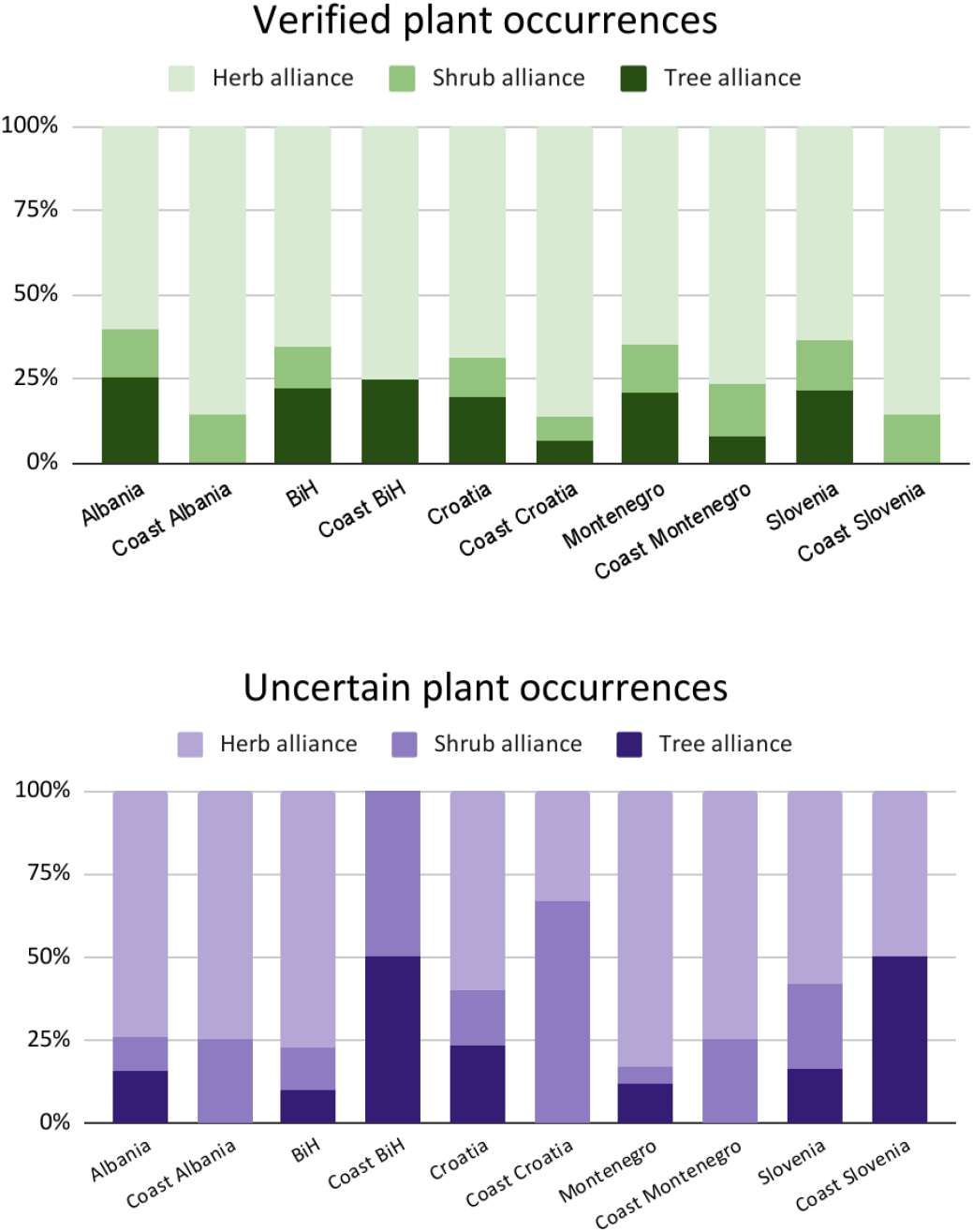
The percentage of verified and uncertain plant alliance occurrences on land and coastland, Albania, Bosnia and Herzegovina (BiH), Croatia, Montenegro and Slovenia

## Conclusions

Database of European Vegetation, Habitats and Flora (floraveg.eu) certainly provided a source of data information on plant alliance occurrences, and on plant alliance missing data, in Albania, Bosnia and Herzegovina, Croatia, Montenegro and Slovenia, neighboring countries in Southeastern Europe. For example, tree alliances were still to be verified on coastland in Bosnia and Herzegovina and Slovenia, and herb alliance on land in Montenegro, Bosnia and Herzegovina and Albania. Differences were found between verified and uncertain plant alliances on land and coastland and in between countries. Overall, herb alliances were found, yet, to be further identified on land in the wild, contributing to characterize the vegetation habitat types in the study region (Preislerová *et al*., 2024).

